# Sensitivity and uncertainty in the Lee-Carter mortality model

**DOI:** 10.1101/2023.01.31.526522

**Authors:** Wenyun Zuo, Anil Damle, Shripad Tuljapurkar

**Affiliations:** Department of Biology, Morrison Institute for Population and Resource Studies, Stanford University, Stanford, CA 94305-5020, USA; Department of Computer Science, Cornell University, Ithaca NY 14853, USA

## Abstract

**BACKGROUND:** The Lee-Carter model (LC) is widely used in research and applications for forecasting age specific mortality, and typically performs well regardless of the uncertainty and often the limited quality of mortality data.

**OBJECTIVE:** Why dose LC perform well regardless of the uncertainty and the limited data quality?

**METHODS:** We analyze the robustness of LC using sensitivity analyses based on matrix perturbation theory, coupled with simulations that examine the effect of unavoidable randomness in mortality data. The combined effects of sensitivity and uncertainty determine the robustness of LC.

**RESULTS:** We find that the sensitivity of LC and the uncertainty of death rates both have nonuniform patterns across ages and years. The sensitivities are small in general, with largest sensitivities at both ends of the period. The uncertainty of death rates are high in young ages (5–19) and old ages (90+) with rising in young ages but dropping in old ages.

**CONCLUSIONS:** The sensitivity and uncertainty analyses indicate that the randomness hardly interrupts the prediction of LC. The divergence of long term prediction in LC is likely due to structural changes in the age-year specific mortality, which is associated with the development of public health policy, medical innovations and chronic disasters.

**CONTRIBUTION:** Our results reveal that LC is robust against random perturbation and sudden short term changes.

## 1 Introduction

The Lee-Carter model (henceforth LC) was developed in 1992 to predict mortality in the US (Lee and Carter, 1992) and has become a widely used tool for mortality forecasting (Wilmoth, 1998; Hollmann, Mulder, and Kallan, 2000; Tuljapurkar, Li, and Boe, 2000; Lee and Miller, 2001; Booth and L. Tickle, 2008; Hyndman and Ullah, 2007; Cairns, Blake, and Dowd, 2008; Girosi and King, 2008). The model is typically fit for each sex to mortality data from a base period, and then used to forecast mortality, e.g. we might use a base period 1961-1990, and forecast mortality for the period 1991-2020. LC (and technical variations mentioned below) typically produces good forecasts in this setting (goodness being assessed in several ways, e.g. Lee and Miller (2001) and Cairns, Blake, Dowd, et al. (2011)).

In at least two respects the success of LC is puzzling. Firstly, why should we expect such a model to successfully predict future mortality? Indeed, at old ages there were fewer people and deaths in the base period relative to the future. So how does the model “know” what will happen at old ages that are poorly represented in the base period? Secondly, why do LC forecasts appear robust to limitations on data quality such as uncertainty in old ages, or any age, mortality rates in any base period, or poor or missing mortality data in developing countries (Keilman, 1997; Girosi and King, 2007)?

To address these questions, we evaluate the robustness of LC when fitted to age-specific mortality rates over any given base period. Here robustness reflects two issues: the sensitivity of LC to changes in the base mortality rates, and the uncertainty that we expect in these rates. The combined effects of sensitivity and uncertainty determine the robustness of this (or any) model. A model is robust when model sensitivity and the uncertainty in the underlying data offset each other (high sensitivity meets low uncertainty or vice versa). On the other hand, we expect poor performance when model sensitivity amplifies uncertainty in the data at some or all ages (high sensitivity meets high uncertainty). The sensitivity of LC is evaluated here for the first time (to our knowledge) by using methods from matrix perturbation theory to compute model sensitivities by age and time. We combine sensitivities with the uncertainty of mortality data (i.e., the variances) of mortality rates to obtain a total measure of model robustness, using numerical simulations to complement our analytical results. We are able to use this approach because LC uses the singular value decomposition (henceforth simply denoted as SVD) method (Lee and Carter, 1992; Tuljapurkar, Li, and Boe, 2000; Lee and Miller, 2001) applied to a matrix of death rates in the base period. In terms of errors in the data, we consider only random errors due to finite population sampling, although our sensitivity measures can be used to examine the effect of other kinds of error.

Our results emphasize the dependence of LC results on the base period used in the forecast and should inform generalizations of the model-fitting approach in LC, such as the functional singular value decomposition (Hyndman and Ullah, 2007), different fitting methods (De Jong and Leonie Tickle, 2006; Brouhns, Denuit, and Vermunt, 2002), and related Bayesian approaches (Girosi and King, 2007).

LC has both age and time dimensions (summarized below). We find that in both dimensions the LC fit by SVD to any recent 50-year base period is remarkably robust. The sensitivities with respect to age-year specific mortality on dominant singular value of the fitted SVD used in LC are small, with the largest sensitivities at the young ages at the start and end of the base period.

Even though the uncertainty of old age mortality rates is high in the early years because of small populations at those ages, this randomness is offset by the low sensitivity at those ages. The year-specific sensitivity on time parameter of LC is low at the middle of the base time period and high near the start and end, as noted long ago by Lee and Carter (1992). While this pattern reflects secular mortality decline, it also follows that LC is less affected by data uncertainty for the years near the middle of the base period. The robustness of LC to uncertainty at old ages and near the middle of the base period contribute to the performance of the method in past decades.

However, we find that LC is more sensitive to mortality rates at the young ages (5–19) as shown in our results. This is important because we expect (Lee and Tuljapurkar, 1997; Zuo et al., 2018) that the peak ages of death will continue shifting to progressively older ages, whereas mortality rates at the young ages continue to decline. The consequence of the latter will be an increase in the uncertainty of young-age mortality rates which will be a challenge to its future performance in low mortality countries.

## 2 Method and results

Here we summarize the essentials of LC and the use of an SVD. LC is applied to a time series of age-specific mortality rates. The original method used central death rates, but here we use age-specific probabilities of death (either choice works). We assume that annual rates are available for *n*_*A*_ single year age classes (with a final open-ended class) for a given base period of *n*_*T*_ years. Write *q*_*at*_ for the 1-year probability of death (central death rate in LC) for age class *a* in calendar year *t*. The mean-centralized natural logarithms of the mortality probabilities (at age *a* and year *t*) are

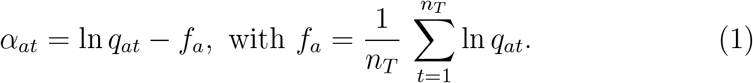

LC uses values of *q*_*at*_ in the base period to fit values of vectors **b** describing the age-profile of mortality change with age-specific components {*b*_*a*_}, and **k** describing the mortality trend over time with year-specific components {*k*_*t*_}, to get the model

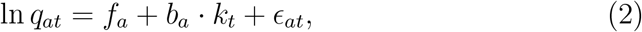

where *ϵ*_*at*_ is a random error. In many cases, the dynamics are well described by assuming that *k*_*t*_ follows a simple random walk with constant drift and diffusion over time, and *f*_*a*_ and *b*_*a*_ are age dependent but time independent.

LC uses an SVD of a matrix **A** whose elements are the base period values of the *α*_*at*_ in equation (1). An SVD of **A** yields singular values and corresponding singular vectors that provide a decomposition of the data matrix. We refer to the fraction of variance in the base period data (over age and time) that is explained by the singular vectors associated with the dominant singular value (*σ*_1_) as the explanation ratio *ϕ*_1_. The value *ϕ*_1_ is large for the US (Lee and Carter, 1992) and for many other countries (Tuljapurkar, Li, and Boe, 2000), so LC approximates the mortality dynamics in terms of *σ*_1_ and the corresponding singular vectors, **u**_1_ with age-specific components {*u*_1*a*_}, and **v**_1_ with year-specific components {*v*_1*t*_}. Accordingly LC sets

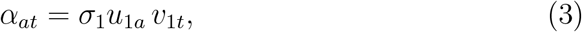

and uses the transformations

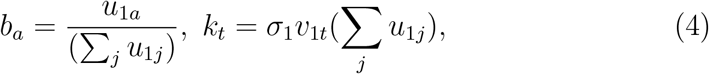

to arrive at the canonical LC model in equation (2).

### Assessing robustness

The combined effects of sensitivity and uncertainty determine the robustness. We examine the sensitivity of LC to changes in the base mortality rates and discuss the uncertainty that we expect in these rates.

To address the effect of systematic errors in the mortality rates, we assess the sensitivity of the dominant singular value *σ*_1_, the age parameter **b** and time parameter **k** to small changes in mortality rates. We use perturbation theory of the SVD that characterizes the behavior of the dominant singular value of the SVD (Weyl, 1912) and singular vectors (Wedin, 1972; G. W. Stewart, 1973; G. Stewart and Sun, 1990). In particular, for the singular values we can use an expansion (Kato, 1976) that is quite accurate provided the perturbation is “small” relative to the separation between singular values and their distance from zero (the larger the separation/distance, the larger the perturbation that can be reasonably handled by the expansion). For the singular vectors we can use results that bound the change in singular subspaces due to a perturbation.

Next, we address the uncertainty of mortality rates (random errors) that occur because mortality is an outcome of a random process. We assess random errors expected in the base data and how these change over time. Then we use simulations and analysis to assess the effect of random errors on *σ*_1_, *ϕ*_1_, *b*_*a*_ and *k*_*t*_.

We illustrate our results using data on the US and Japan. We find (but do not report) qualitatively similar results for other countries. Annual single-year probabilities of death from 1950 to 2014 were obtained from the Human Mortality Database (HMD) (University of California, Berkeley (USA) and Max Planck Institute for Demographic Research (Germany), 2018). For our purposes the HMD death rates at ages over 95 years may be substantially affected by the use of a model, so we present results only for ages through 95 with a single open age class at ages 95+. We fit an SVD (i.e., LC) using a 50 year base period: the earliest base period is 1950 to 1999, and the most recent is 1965 to 2014. Where we consider only a single base period, we report results for the earliest base period, since results for other base periods are qualitatively similar.

To illustrate the age and time dimensions of LC, we compute the dominant singular vectors **u**_1_ and **v**_1_ for all base periods. Fig. 1 plots values of **u**_1_ versus age *a* for the US (top left) and Japan (top right); and corresponding plots for **v**_1_ versus time *t* (lower panel). The black curves are for 1950-1999, while the gray curves represent base time periods shifted in increments of 5 years (a similar figure for **b** and **k** is Fig. A.1). For all base periods, the curves have a general pattern that is familiar to anyone who has used LC. Notice that the Japanese data generates noticeable differences in **u**_1_ for different base periods and smaller differences in the **v**_1_, a pattern that is reversed for the US. Over those four base periods the dominant singular values are in the range 14 *−* 16 for the US and 25 *−* 34 for Japan. The explanation ratio (*ϕ*_1_) is about 0.95 for the US and 0.97 for Japan for all four base periods (differs in the third decimal digit). The large dominant singular value for the US and Japan is present in different base periods, which can be explained by the sensitivity of the dominant singular value (details in the section of “Sensitivity of the dominant singular value”).

**Figure 1.**
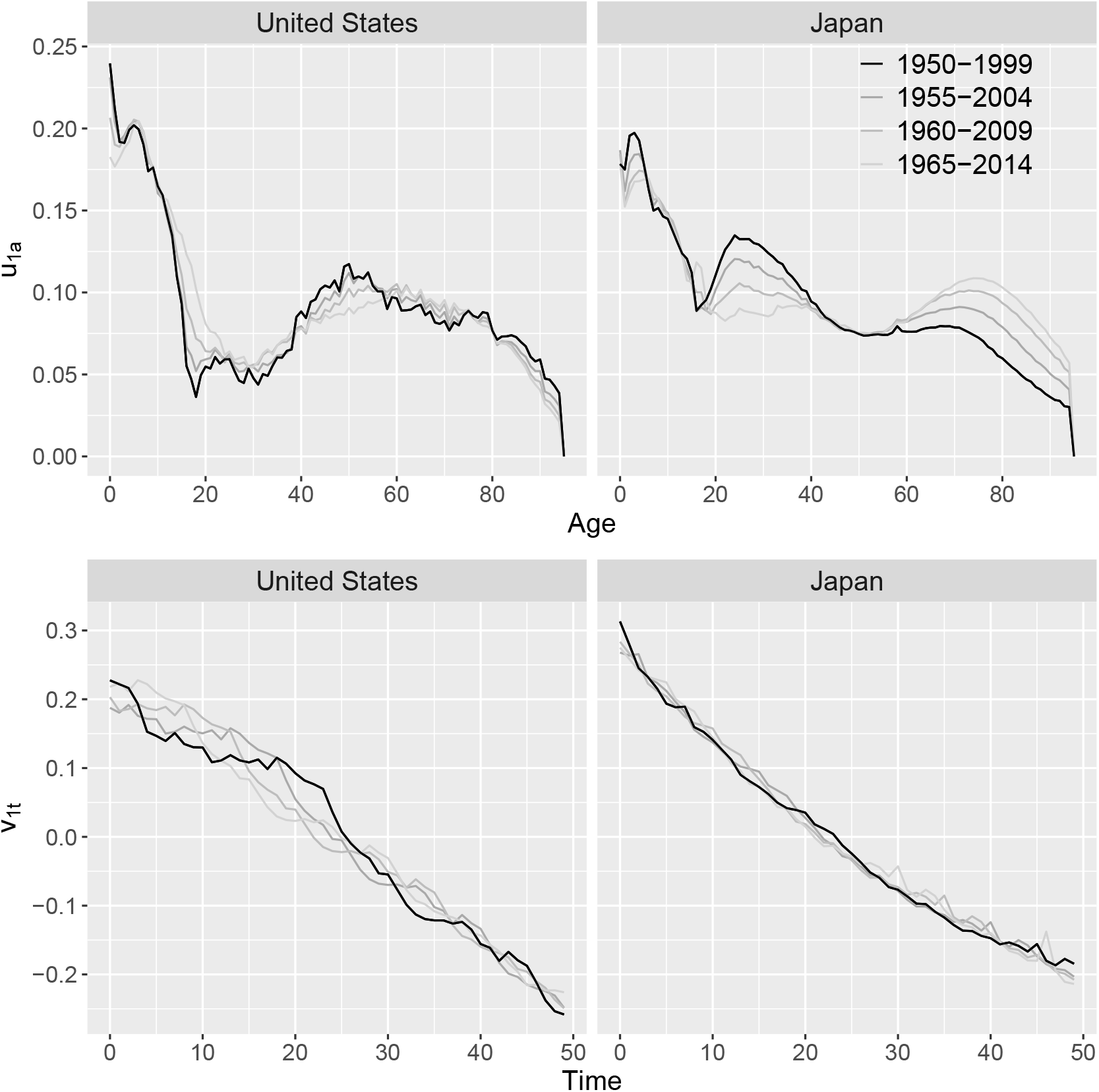
Singular vectors **u**_1_ with age-specific components *u*_1*a*_, upper panel, and **v**_1_ with year-specific components *v*_1*t*_, lower panel for US (on left) and Japan (on right). They determine sensitivities of dominant singular as in equation (5). Different base time periods from 1950-1999 onward, advancing every 5 years. The black curves are 1950-1999; the lighter the gray, the later the period. See a similar figure for LC parameters, age-profile of mortality change **b** and the mortality trend over time **k**, is in Fig. A.1.

### The effect of small changes in mortality

We start with the effect of small mortality changes (here, also called perturbations), so that **A** is replaced by a new matrix **Â** = **A** + **E**, where **E** has elements {*e*_*at*_} and is considered small relative to **A**. In place of the original dominant singular value and vectors *σ*_1_, **u**_1_, **v**_1_, an SVD of **Â** yields new dominant singular value and vectors that we call 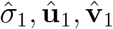. In terms of LC, equation (4), this corresponds to changes from the original **b, k** to 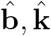.

### Sensitivity of the dominant singular value

Perturbation theory for singular values can be assessed via expansions (Kato, 1976) and bounds (Weyl, 1912) and the use of said expansions shows the sensitivity of *σ*_1_ to the logarithm of death rate at (*a, t*) is

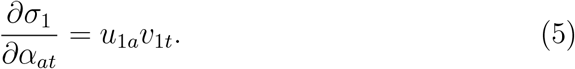

Details of the derivation of equation (5) are in the appendix section A.3.

Importantly, a small change in single mortality *q*_*at*_, written as ln *q*_*at*_ + *δ*, leads to multiple changes in **A**. Equation (1) indicates that ln *q*_*at*_ + *δ* = *α*_*at*_ + *δ − δ/n*_*T*_, where *n*_*T*_ is the length of the base period. At all other *i ≠ t*, the change in the logarithm is *α*_*ai*_ *−δ/n*_*T*_. Therefore, the change in the dominant singular value with respect to the the logarithm mortality matrix is the sum of all the changes in **A**.

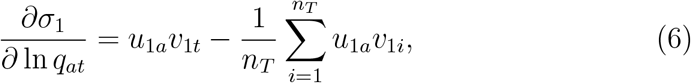

which gives the sensitivity of the dominant singular value, denoted by 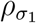, as

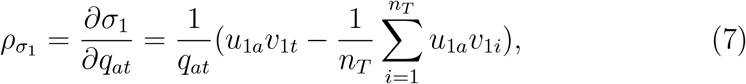

Since 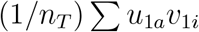 is small for long base periods we can approximate equation (7) as

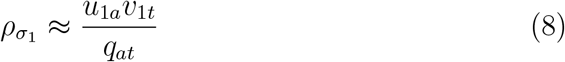

The patterns shown in Fig. 1 illustrate this sensitivity of *σ*_1_ over the four base periods. As expected from equation (8) the sensitivity of dominant singular value to the changes in mortality in all ages is lowest near the middle of the base period, where *v*_1*t*_ *≈* 0, but larger and positive at the start of the period, and larger and negative towards the end. In general, small values of *u*_1*a*_ at high ages, regardless of the year *t*, imply a low age-specific sensitivity at old ages, and offset the large uncertainty in death rates at these ages. This is one reason why LC worked so well in the past, and explains how LC appears to “know” what will happen at old ages that are poorly represented in the base period.

As the base period advances, in the US the values of *u*_1*a*_ decrease at old ages, and there are changes in *v*_1*t*_ but with no clear trend. In contrast, in Japan the values of *u*_1*a*_ increase at old ages, especially around age 75 yrs, but there are only modest changes in *v*_1*t*_. Given that old-age mortality fell over time, equation (8) suggests a modest positive change at *σ*_1_ due to old ages for the US and a large but negative change at *σ*_1_ for Japan. Therefore, we expect and find that *σ*_1_ for the US rose from 14.0 to 16.2 but for Japan fell from 34.4 to 25.0 over the four base periods shown (Table A.1 in Appendix). However, note the higher age specific sensitivity of the dominant singular value at young ages (under 10), especially near the start and end of the base period. This may be a challenge in the future as we discuss later.

The explanation ratio *ϕ*_1_ that describes the relative importance of the dominant singular value characterizes the goodness of fit of LC to the mortality. As many of the singular values are less well separated from others than the dominant one, the perturbation expansions are of little use here. Formally, Mirsky’s Theorem (“Symmetric gauge functions and unitarily invariant norms” 1960) can be used to bound the net change in singular values in terms of the size of the perturbation allowing for some analysis of *ϕ*_1_. However, given the relative dominance of *σ*_1_ for the base periods used here (resulting in *ϕ*_1_ near 0.95 for the US and 0.97 for Japan) the explanation ratio does not change significantly over the four base periods even though *σ*_1_ changes for the base periods.

### Sensitivity of age and time parameters

The relationship between LC and the SVD, equation (4), shows that sensitivity of the LC age parameter **b** and time parameter **k** can be understood by characterizing the changes in **u**_1_, **v**_1_ and *σ*_1_. As with the singular values, it is possible to develop first order expansions for the singular vectors. However, for our purposes upper bounds on the change in singular vectors suffice to illustrate the key points. While we defer a full treatment to Appendix A.4, we provide an informal statement of the results here. As before, consider **Â** = **A** + **E**. If Δ**u** and Δ**v** denote the change in left and right dominant singular vector separately, where Δ**u** = **û**_1_ *−***u**_1_ and 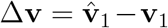 and **û**_1_ and 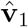 are the left and right dominant singular vector of **Â** then for sufficiently small *E* we have that

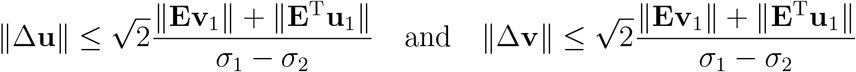

In other words, the change in the singular vectors is strictly bounded by how well the perturbation aligns with **u**_1_ and **v**_1_ (which is bounded by the size of **E** since the singular vectors have unit norm) divided by the separation between the first and second singular values (a quantity we expect to be large given the explanation ratios close to 1).

To consider how perturbations to the singular vectors affect the LC model let 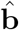 and 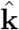 be the age and time parameters of perturbed matrix **Â**. Equation 4 yields

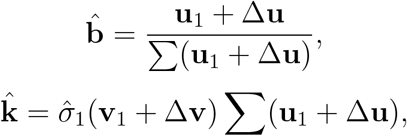

and we define the difference in age vector as

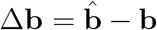

and the difference in the time vector as

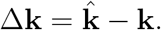

Similarly, we define the age-year specific sensitivity of **b**, *ρ*_*b*_, as

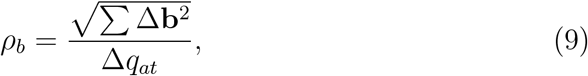

and the Euclidean distance between **b** and 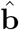 per unit change of age-year specific mortality and the age-year specific sensitivity of **k**, *ρ*_*k*_, as

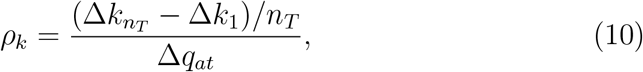

the changes in slopes of year vector over time between **k** and 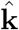 per unit change of age-year specific mortality. These sensitivities of **b** and **k** turn out to be small, with largest sensitivities at two ends of the base period and young ages, which is similar to the sensitivity of dominant singular value. Furthermore, age parameter **b** shows an intriguing pattern. The perturbation at a given age, *τ*, regardless of the year (except for years in the middle of the base period) has its largest impact on the *τ*_th_ component in **b**. The sensitivity of **b** in the middle years of the base period is close to zero.

### Uncertainty in mortality rates

We now assess the effect on LC of unavoidable random errors in mortality rate, given that mortality rate is an outcome of a random process. What random errors, i.e., variances, should we expect? To be clear, the variance here is the potential random errors for each single age-year specific mortality rate, different from the mortality variance-covariance matrix used in the previous section.

Assume that we know accurately the population at risk, *n*_*at*_, and that deaths follow a Poisson distribution (Brillinger, 1986; Brouhns, Denuit, and Vermunt, 2002). The death rate is a random variable, say *Q*_*at*_ whose mean is the base rate *q*_*at*_ and whose variance is

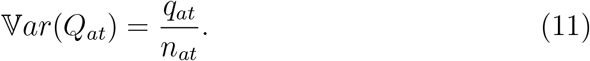

Hence the variance of ln *q*_*at*_ is approximately

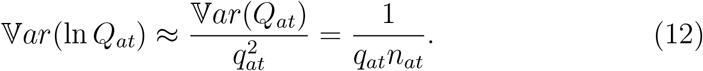

What matters to LC are these logarithmic variances. The rates in LC are actually random variables of the form *α*_*at*_ + *e*_*at*_, with 𝔼(*e*_*at*_) = 0 and

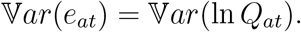

Equation (12) shows that an increase in variance at age *a* in year *t* is produced by small number of death which is contributed by either a small death rate *q*_*at*_ and/or a small population at risk *n*_*at*_.

For the dominant singular value, we can combine this variance with the earlier perturbation results to get the approximation

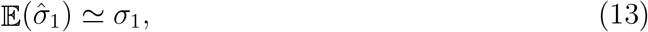

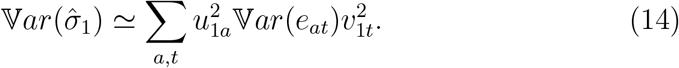

Thus we expect that the dominant singular value should be least robust to death rates at ages where there are relatively high values of both the uncertainty of death rates and the sensitivity of the dominant singular value 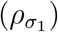, and vice versa.

More generally, we can use simulation to generate sets of “base” random death rates by setting

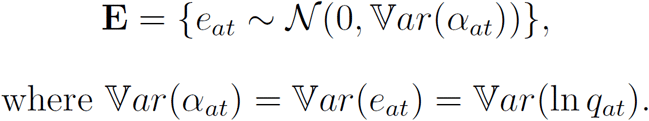

The choice of a normal distribution here is made for speed and convenience; trials with skewed distributions such as the Poisson did not affect our conclusions. Using the base period 1950-1999, we used simulations to examine variation in the dominant singular value and the explanation ratio. Every randomly perturbed matrix **Â** = **A** + **E** has a dominant singular value 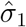 and an explanation ratio 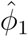. We computed these by doing an SVD on each of 1000 randomly perturbed matrices (More details in Appendix A.2).

The simulations agree with the analytical result of equation (13), in that the average 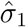 from simulation is only 0.001 larger than the *σ*_1_ from base period matrix for both the US and Japan. In equation (14), use the values of the random variation of age-specific annual mortality rate in Fig. 2 together with the patterns of base period dominant singular vectors in Fig. 1 to see that the variance in the dominant singular value is small. The simulations agree, with a variance of 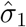 near 10^*−*4^ for the US and near 10^*−*3^ for Japan. Equation (14) also shows that the uncertainty of age-specific mortality does not contributed to the variance of the dominant singular value uniformly. The simulations reveal that 57.2% of this variance contributed by ages 5–19 for the US and 54.2% for Japan. For both the US and Japan, there are little covariance between age groups.

**Figure 2.**
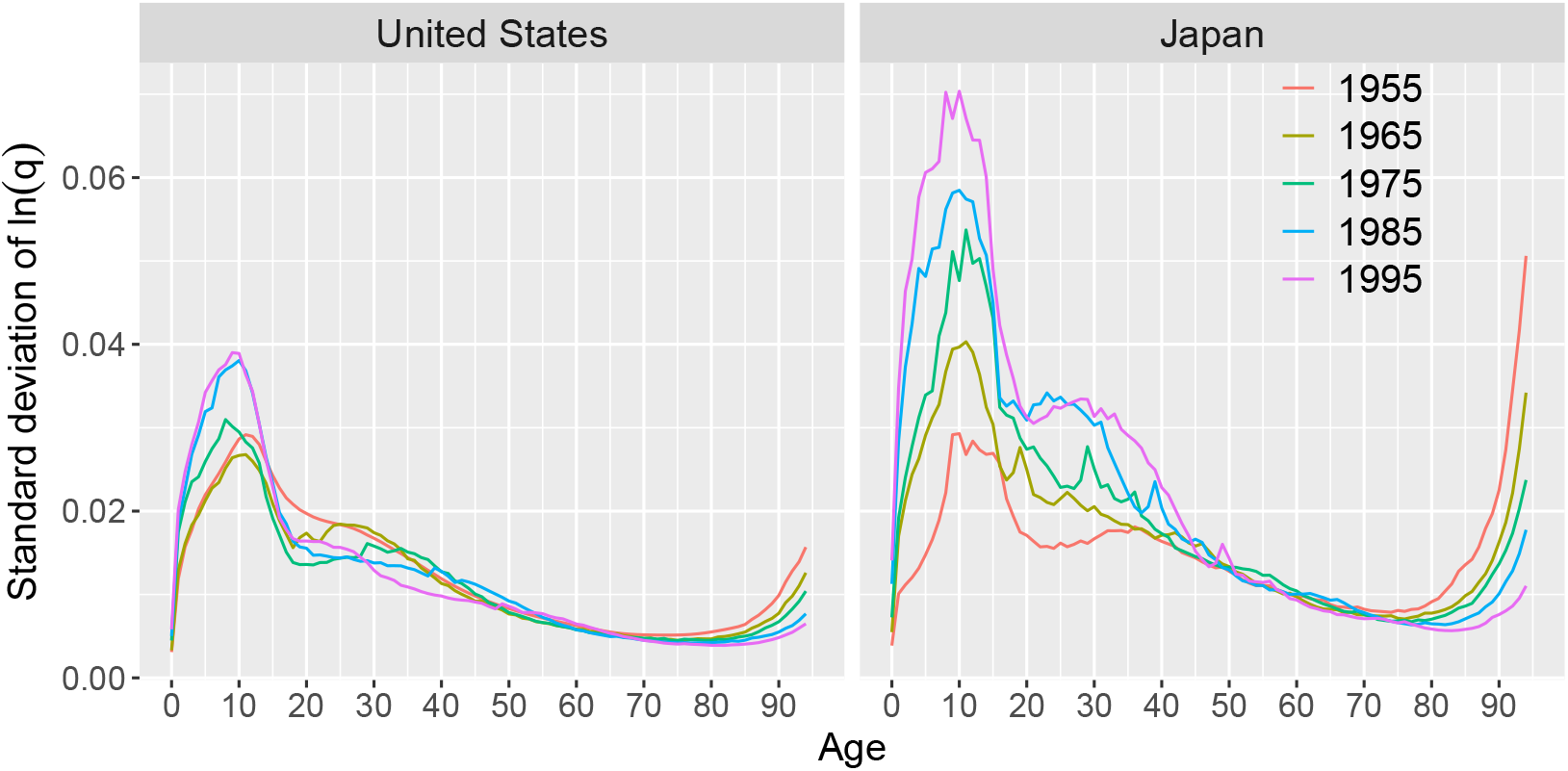
Uncertainty of age-specific annual mortality rate over time. Standard deviation (SD) of the annual ln(*q*_*at*_) versus age plotted for every tenth year from 1955 to 1995 for US and Japan.

The simulations also reveal what happens to the explanation ratio. We find that the explanation ratio decreases based on the simulations of the random variation in mortality, and the variance of the explanation ratio from the simulated matrices is also small (Fig. 4). For the US, random variation in mortality decreases the explanation ratio by 0.4% to 94.2%, and for Japan decreases by 0.2% to 97.2%. How could the small increase (0.001) of dominant singular value cause this decrease of *ϕ*_1_? Our simulations show that random variation in mortality changes (increases) little in the dominant singular value, *σ*_1_, but changes (increases) more in other singular values. The changes in the explanation ratio could be a useful diagnostic for extensions to LC.

The uncertainty in mortality rate changes over time, as shown by the standard deviation of the natural logarithmic death rates for years 1955-1995 in Fig. 2 for the US and Japan. The largest uncertainties are at ages 5–19 and 90+; in both age ranges there is a relatively small number of deaths but for quite different reasons. Young ages have very low mortality rates in developed countries and a large population at risk, while the opposite is true at old ages. Using age 10 as an example, from 1950 to 1999 the mortality probability per 10,000 fell in the US by 250% (from 4.4 to 1.7) and in Japan by 490% (from 6.9 to 1.4). Over the same years, the exposure in that age class rose by 40% in the US but fell by 15% in Japan. In contrast, at age 94, in both countries death rates fell by much smaller proportions but exposures increased by far larger proportions. These trends caused the uncertainty of mortality at both ages to change substantially over time (Fig. 3).

**Figure 3.**
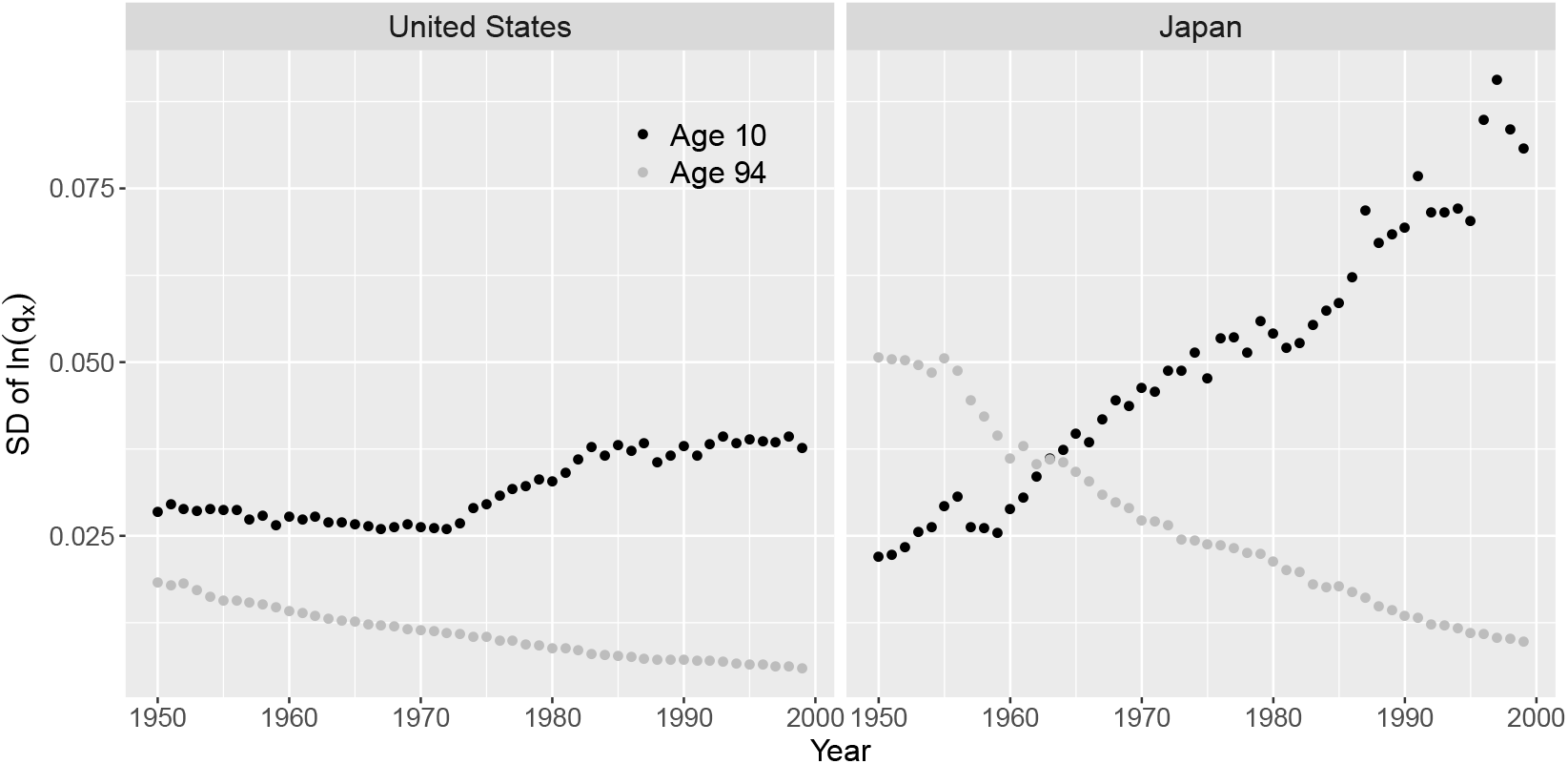
The uncertainty of mortality, standard deviation (SD) of ln *q*_*at*_, at age 10 and 94. Over time SD of ln *q*_*at*_ increase at age 10, but decrease at age 94. HMD data of US and Japan as examples presented here.

**Figure 4.**
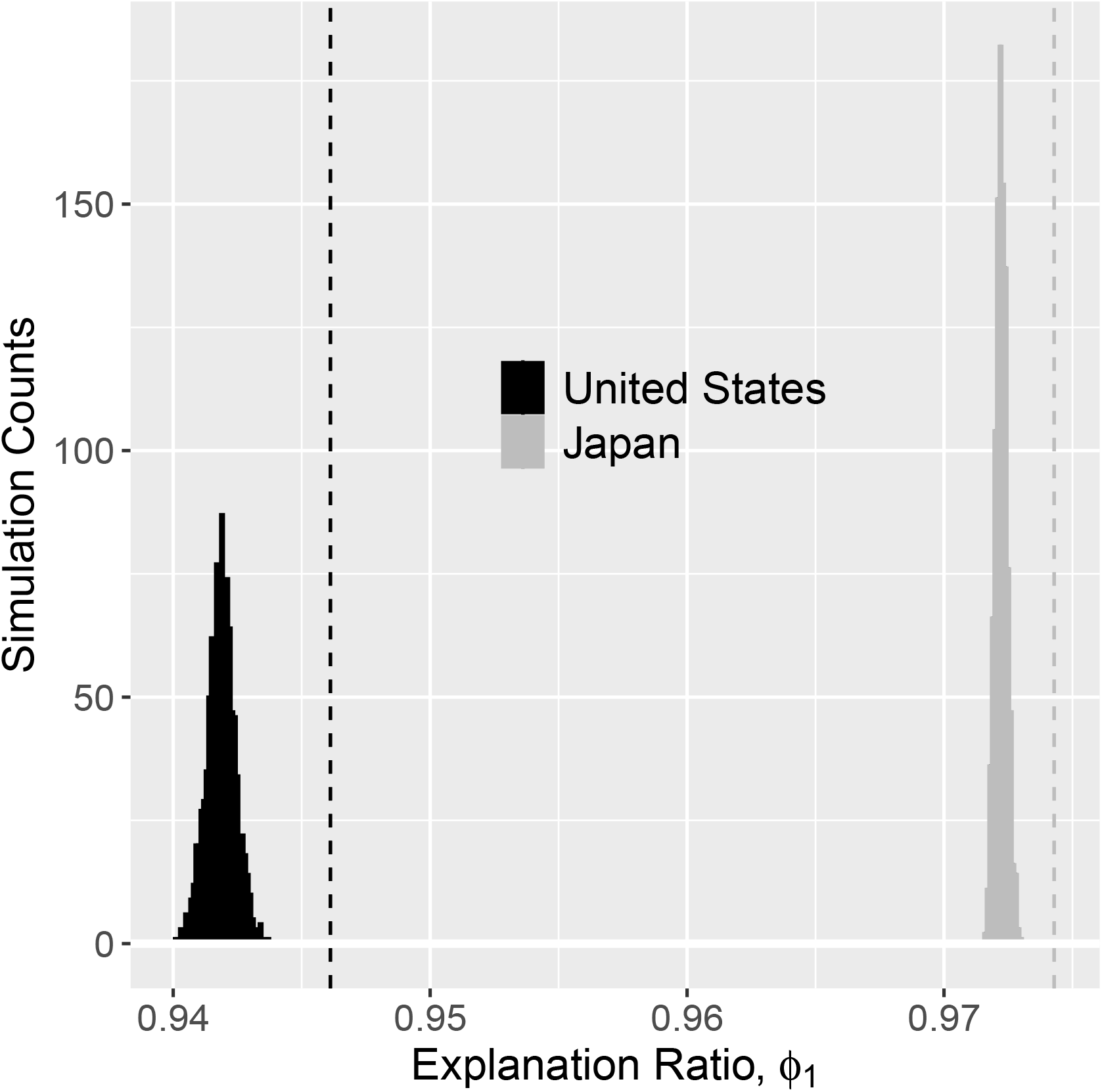
Random variability in death rates affects the explanation ratio, fraction of data variance explained by the dominant singular value for original and perturbed matrices. Dashes lines are explanation ratios of original matrices for US (black) and Japan (gray). Histogram are the explanation ratios from randomly perturbed matrices for US (black) and Japan (gray).

## 3 Discussion

Why does LC perform so well in forecasting given the variability of age-year specific mortality (*q*_*at*_) in the past? We have shown that the answer lies in two aspects associated with LC, the sensitivity of LC parameters to changes in *q*_*at*_ and the uncertainty of *q*_*at*_. The robustness of LC is determined by the robustness of the LC age parameter (**b**) and time parameter (**k**). The robustness is the combined effect of sensitivity and uncertainty, which means that the robustness of **b** is the entry-specific product of the sensitivity of **b** (*ρ*_*b*_) and uncertainty of *q*_*at*_ and the robustness of **k** is the entry-specific product of the sensitivity of **k** (*ρ*_*k*_) and uncertainty of *q*_*at*_.

We found that the sensitivities of **b** and **k** to changes in *q*_*at*_ are not evenly distributed (equations 9 and 10), which indicates that LC is sensitive to mortality changes in some years and ages more than others. The magnitude of the unevenness is enormous. Here we use the US for the base period 1950 to 1999 as an example to provide an intuitive view. The largest sensitivity of **b** in the US is to the mortality of age 10 in 1999 with *ρ*_*b*_ = 13 which is about 10^5^ times larger than the smallest *ρ*_*b*_ at age 0 in 1976 with *ρ*_*b*_ = 6 *×* 10^*−*5^. Similarly, the largest sensitivity of **k** in the US is to the mortality of age 10 in 1999 with *ρ*_*k*_ = 205 which is about 10^6^ times larger than the smallest *ρ*_*k*_ at age 0 in 1976 with *ρ*_*b*_ = *−*7.5 *×* 10^*−*5^.

We also found the uncertainty of *q*_*at*_ is not uniformly distributed across entries, being associated to the number of deaths at a given age in a given year. A small number of deaths means large uncertainty, equation (12). There are two paths which lead to a small number of deaths, small population at risk or small mortality. Both the US and Japan indicate that old age groups (90+) and young age groups (5–19) have large uncertainty of *q*_*at*_ in the past. Old age groups have large uncertainty due to small population at risk. Fortunately the old age population at risk increases over time (Zuo et al., 2018), so that the uncertainty decreases. Young age groups have large uncertainty due to small mortality rate. With improvements of economy, medical care and public health, *q*_*at*_ at young age groups is expected to continuously decrease over time, which unfortunately will further increase the uncertainty of *q*_*at*_ at young age groups (Fig. 3).

LC is clearly quite robust regardless of the significantly uneven effects on **b, k** and the uncertainty of *q*_*at*_ across ages and years to changes of *q*_*at*_. Still using the US for the base period 1950 to 1999, the largest sensitivities of **b** (*ρ*_*b*_ = 13) and **k** (*ρ*_*k*_ = 205) both are at age 10 in 1999. The mortality of age 10 in 1999 is only 1.5 *×* 10^*−*4^. Therefore, even a 100% change in mortality hardly impacts the age vector **b** in LC. Similarly, a 100% change in mortality leads to a change of 0.03 in the slope of year vector, which means that prediction of future *k*_*t*_ is stable.

LC has small number of parameters 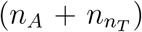 and large number of observed data points 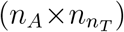. Moreover, the relative robustness of singular values and vectors for well separated singular values (as is common here) ensures that LC is an effective forecasting method even in the presence of uncertainty in the underlying data.

In the future (if not already), mortality will have higher uncertainty in young age groups (5–19) than old age groups (90+) as mentioned above. The increasing uncertainty in young ages has impact on the life expectancy forecasting, because life expectancy is more sensitive to young ages than to old ages (Vaupel, 1986).

LC is robust against random perturbation caused by unavoidable sampling errors from the uncertainty of mortality rates and some sudden changes in mortality for a given year such as the mortality rate spike in Japan in 2011 due to the tsunami. Even though the increasing uncertainties of *q*_*at*_ at young ages affect the robustness of LC, the magnitude of the effect should be small due to the limited changes in mortality rate (a 10 times increase of the mortality of age 10 in 1997 is only 1.5 *×* 10^*−*3^) and the absolute values of *ρ*_*b*_ and *ρ*_*k*_. But one should ask if LC is so robust why is there a significant divergence of long term prediction in the LC? The divergence of long term prediction in the LC is not due to mortality rates sampling errors or sudden changes, but it most likely due to structural changes in the LC age parameter **b**, which is associated with the development of public health policy, medical innovations and chronic disasters. Therefore future improvement on LC should focus on how to predict the structure change of age parameter **b**.

## Appendix

### A.1 General features of SVD

An SVD of matrix **A** yields singular values *σ*_1_ *≥ σ*_2_ *≥* … *≥ σ*_*i*_ *≥* 0, for *i* = 1, …, min(*n*_*A*_, *n*_*T*_) with corresponding left singular vectors **u**_*i*_ (with components *u*_*ia*_, *a* = 1, …, *n*_*A*_) and right singular vectors **v**_*i*_ (with components *v*_*it*_, *t* = 1, …, *n*_*T*_). These vectors provide a complete decomposition of the data,

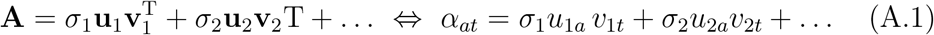

The left and right singular vectors are orthonormal. Each singular vector has unit Euclidean length,

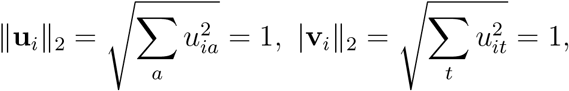

and all the left (respectively, right) singular vectors are perpendicular to each other, i.e.,

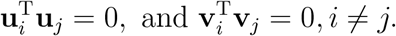

The centering in text equation (1) implies that for every age *a*,

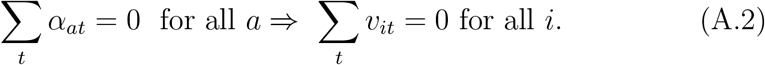

The singular value *σ*_*i*_ measures the fraction of total variance 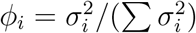 explained by **u**_*i*_ in the age dimension, or **v**_*i*_ in time. Here we will assume *σ*_1_ *> σ*_2_ (as is quite natural in practice), so the vectors **u**_1_, **v**_1_ represent the directions explaining the largest fraction of variance. The US data (Lee and Carter, 1992) have a first singular value that is significantly larger than the rest (as do most mortality matrices (Tuljapurkar, Li, and Boe, 2000)). The larger the proportion of variance explained by the dominant singular value 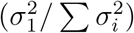, the better is the fit of LC to death rates in the base period.

### A.2 Simulation

Each simulation discussed here has a set of 1000 perturbed matrices **Â** _*i*_, where **Â** _*i*_ = **A** + **E**_*i*_ and we can use simulation to generate sets of “base” random death rates by setting

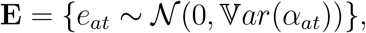

The choice of a normal distribution here is made for speed and convenience; trials with skewed distributions such as the Poisson did not affect our conclusions. All the statistics for each simulation are calculated from 1000 perturbed matrices **Â**_*i*_. All population exposed to risk recorded as 0 are deliberately set to 1 in order to avoid infinite value in the calculation.

### A.3 The change of dominant singular values due to perturbation: two methods

#### A.3.1 Perturbation theory for the SVD

For small perturbations, standard extensions of results from Kato (1976) shows that

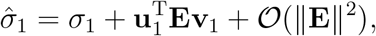

where ∥**E**∥ is the largest singular value of **E**, and 𝒪 (∥ **E** ∥^2^) is at most some constant times ∥**E**∥ ^2^. Note that when dealing with singular values near zero special care must be taken to form viable expansions (G. Stewart, 1984). However, as we are always considering *σ*_1_ the standard formula suffice.

From these expansions we see that the change in the dominant singular value Δ*σ*_1_ can be written as

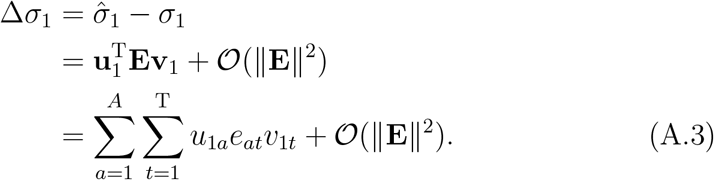

Since *e*_*at*_ is the perturbation of *α*_*at*_, we can write a partial derivative of the dominant singular value *σ*_1_ respect to *α*_*at*_, as

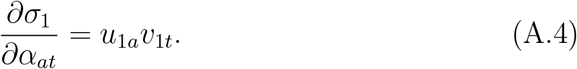

#### A.3.2 Derivation of the singular value expansion

For completeness, we provide a simple (albeit slightly informal) derivation of the expansion used. A full version may be adapted from Kato (1976), see also G. Stewart (1984). As a starting point we first consider the sensitivity of a simple eigenvalue of a symmetric matrix (as singular values can be connected to eigenvalues of symmetric matrices). Specifically, given a symmetric matrix **S** and small symmetric perturbation Δ**M** the eigenvlaues and eigenvectors of **S** + Δ**M** satisfy

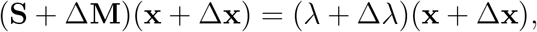

where *λ* and **x** are an eigenvalue/vector pair for a simple eigenvalue. Dropping second order terms (i.e., those containing two perturbations) and using the relation **Sx** = *λ***x** we see that

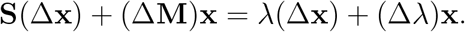

Multiplying both sides by **x**^T^ and using the facts that **x**^T^**S** = *λ***x**^T^ and **x**^T^**x** = 1 we conclude that

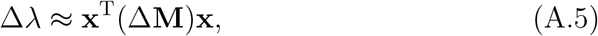

which is accurate to first order.

To extend this result to singular values we may use that if **A** has (reduced) SVD **A** = **UΣV**^T^ then

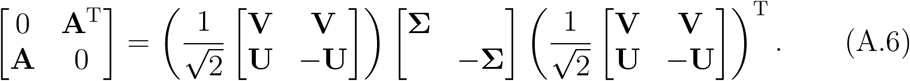

In other words, we can relate the singular values of **A** to those of a symmetric matrix. Assuming *σ*_1_ *> σ*_2_ we can immediately apply Eq. A.7 to the corresponding eigenvalue of

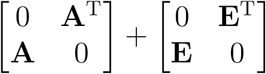

and conclude that

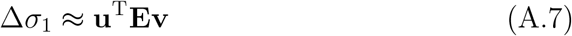

and the expression is accurate to first order. Since **E** is the perturbation matrix, we can write a partial derivative of the dominant singular value *σ*_1_ respect to *α*_*at*_, as

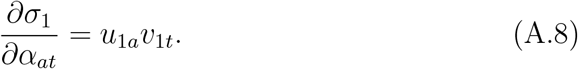

Note that this expression can be derived independently by directly differentiating the eigenvalue/vector equation for a symmetric matrix (see, e.g., Golub and Van Loan (2013)) and then using the same relation for the SVD as above.

#### A.4 The change of dominant singular vectors due to perturbation

Developing explicit expressions and expansions for singular vectors is more complicated than for singular values. However, it is still quite possible to develop strict upper bounds on the changes in singular vectors with respect to a perturbation (G. W. Stewart, 1973; G. Stewart and Sun, 1990). For completeness we provide results here (albeit without proofs). Once again we start with a bound for the changes in the eigenvector of a symmetric matrix associated with a simple eigenvalue and then extend them using Eq. A.6 to apply them to singular vectors.

Given a symmetric matrix **S** and symmetric perturbation Δ**M**. If *λ*_1_ is the algebraically largest eigenvalue of **S** and **x** is the associated eigenvector, classical results^1^ by Davis and Kahan (1970) show that if ∥Δ**M**∥ *<* (*λ*_1_ −*λ*_2_)*/*5 then there exists a Δ**x** satisfying

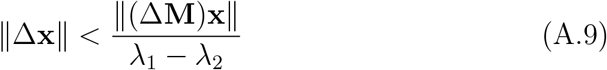

such that **x** + Δ**x** is an eigenvector associated with the algebraically largest eigenvalue of **S** +Δ**M**. Applying this bound to Eq. A.6 immedietly yields the bounds

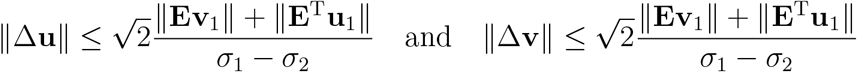

for a perturbation **E**.

**Table A.1:**
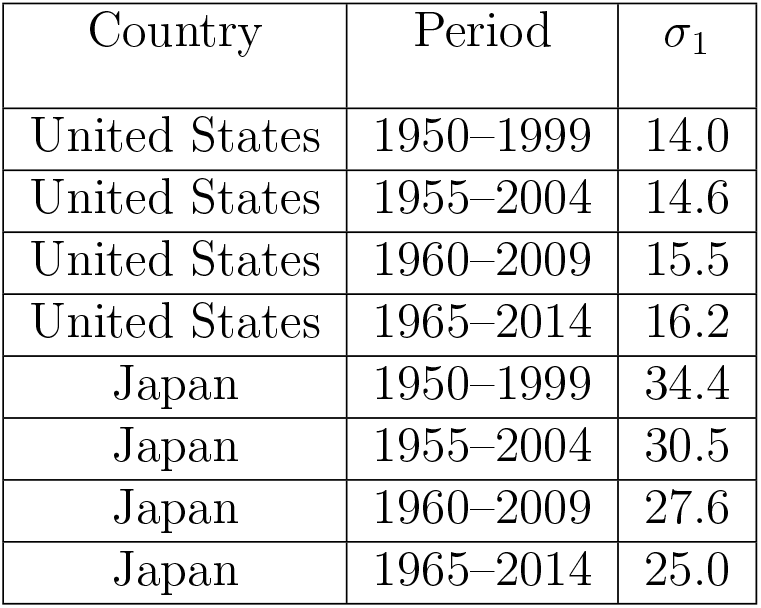
Dominant singular values in different base periods for United States and Japan.

| Country | Period | $\sigma_1$ |
| --- | --- | --- |
| United States | 1950–1999 | 14.0 |
| United States | 1955–2004 | 14.6 |
| United States | 1960–2009 | 15.5 |
| United States | 1965–2014 | 16.2 |
| Japan | 1950–1999 | 34.4 |
| Japan | 1955–2004 | 30.5 |
| Japan | 1960–2009 | 27.6 |
| Japan | 1965–2014 | 25.0 |

**Figure A.1:**
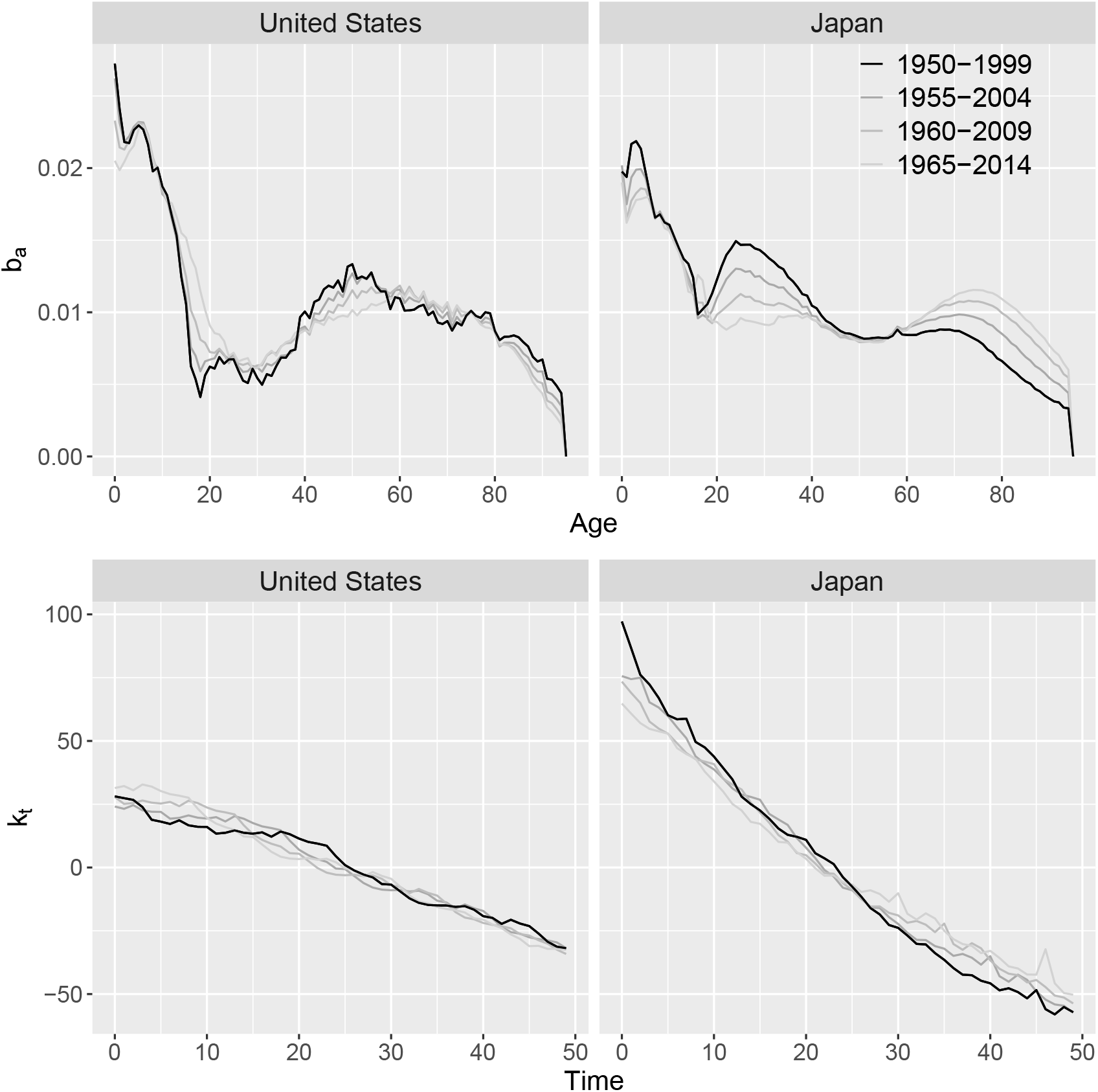
Age-profile of mortality change **b** with component *b*_*a*_, upper panel, and the mortality trend over time **k** with component *k*_*t*_, lower panel for US (on left) and Japan (on right). Different base time periods from 1950-1999 onward, advancing every 5 years. The black curves are 1950-1999; the lighter the gray, the later the period.

**Figure A.2:**
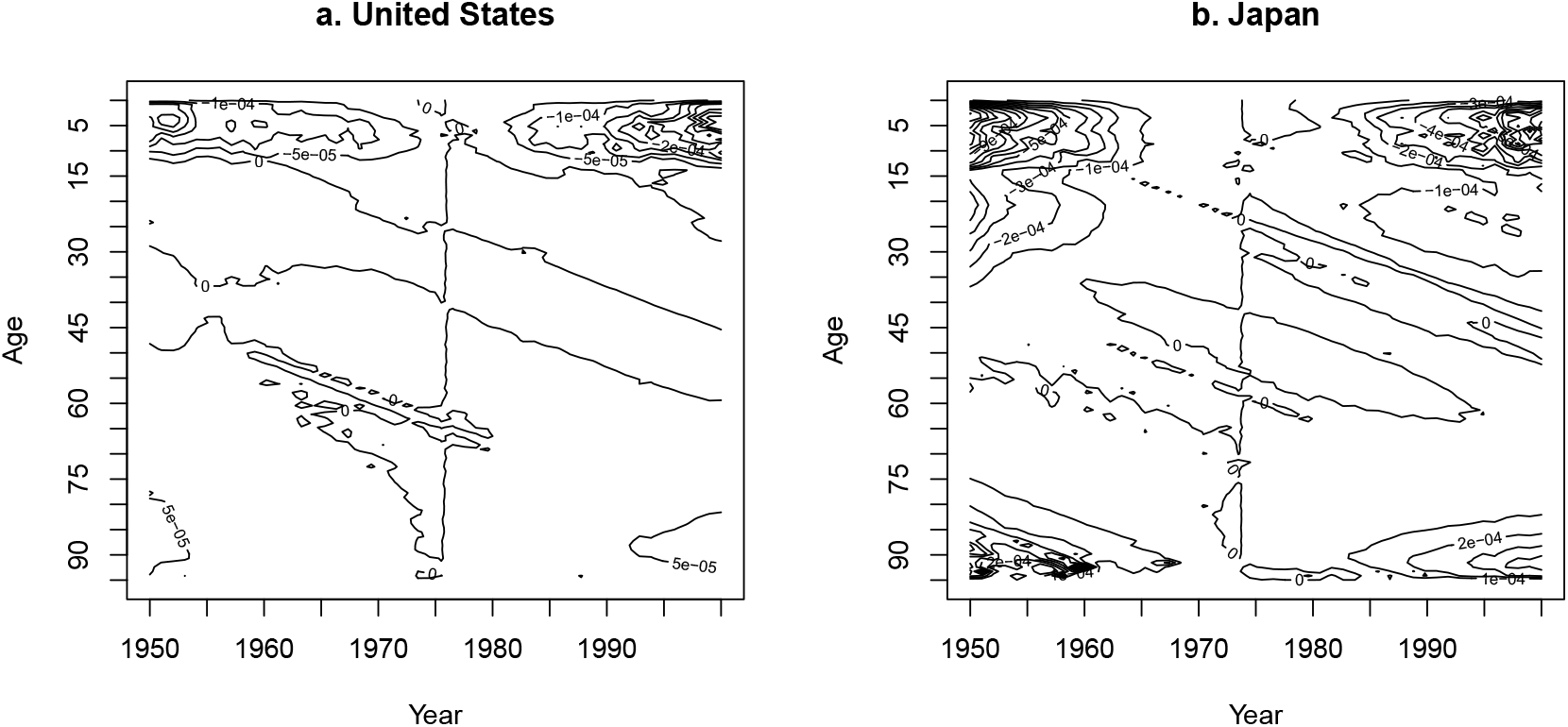
Impact on dominant singular value by increasing 1 standard deviation of the age-year-specific mortality, *q*_*at*_.

## Footnotes

1 Here we have specialized the bounds to the largest eigenvalue; they are far more general.

